# Machine Learning for Missing Data Imputation in Alzheimer’s Research: Predicting Medial Temporal Lobe Flexibility

**DOI:** 10.1101/2025.05.22.655574

**Authors:** Soodeh Moallemian, Abolfazl Saghafi, Rutvik Deshpande, Jose M. Perez, Miray Budak, Bernadette A. Fausto, Fanny Elahi, Mark A. Gluck

## Abstract

**BACKGROUND:** Alzheimer’s disease (AD) begins years before symptoms appear, making early detection essential. The medial temporal lobe (MTL) is one of the earliest regions affected, and its network flexibility, a dynamic measure of brain connectivity, may serve as a sensitive biomarker of early decline. Cognitive (acquisition, generalization), genetic (APOE, ABCA7), and biochemical (P-tau217) markers may predict MTL dynamic flexibility. Given the high rate of missing data in AD research, this study uses machine learning with advanced imputation methods to predict MTL dynamic flexibility from multimodal predictors in an aging cohort.

**METHODS:** In an ongoing study at Rutgers’s Aging and Brain Health Alliance, data from 656 participants are utilized, including cognitive assessments, genetic and blood-derived biomarkers, and demographics. Due to MRI-related constraints, only 34.15% of participants had measurable MTL dynamic flexibility from resting-state fMRI. To estimate MTL dynamic flexibility from available data, we evaluated four missing data handling methods (case deletion, MICE, MissForest, and GAIN), and trained five regression models: Ridge, k-NN, SVR, regression trees (bagging, random forest, boosting), and ANN. Hyperparameters were optimized via grid search with 3-fold cross-validation. Model performance was assessed using mean absolute error (MAE), root mean squared error (RMSE), and runtime through 5-fold cross-validation repeated 25 times to ensure robustness in clinical data settings.

**RESULTS:** A total of 1,866 missing values (25.86%) were identified in the dataset, with only 42 complete cases (6.40%) remaining after listwise deletion, highlighting the need for effective imputation. In the initial analysis using only complete cases, support vector regression (SVR) achieved the lowest mean absolute error (MAE = 0.184), though overall performance was limited due to small sample size. In the second phase, three imputation techniques were applied, significantly improving model accuracy. MissForest combined with Random Forest produced the best results (MAE = 0.083), representing a 54.7% improvement over case deletion. Statistical analysis confirmed significant differences in performance across imputation methods (p < 0.001), with MissForest outperforming GAIN and MICE. GAIN was the fastest imputation method.

**DISCUSSION:** The findings underscore the importance of using robust imputation strategies to maximize data utility and model reliability in studies with high missingness. Further research is needed, particularly incorporating additional neuroimaging measures, to localize the brain regions most affected by biomarker-driven changes and to refine predictive models for clinical applications.

## 2. Introduction

Alzheimer’s disease (AD), the most common form of dementia, is characterized by progressive cognitive decline and widespread neurodegeneration [1]. Pathological changes associated with AD can begin decades before the onset of clinical symptoms [2], underscoring the importance of early detection for the development of effective diagnostic and therapeutic strategies, particularly in the absence of a definitive cure. Integrating multimodal predictors provides a more comprehensive and potentially more accurate means of detecting early signs of AD by leveraging complementary information from cognitive, genetic, and biochemical domains.

### MTL dynamic flexibility

The medial temporal lobe (MTL), particularly subregions such as the transentorhinal cortex and hippocampus, is among the earliest brain regions to undergo neurodegeneration in AD [3,4].

Functional connectivity, defined as the temporal correlation of neural activity between brain regions during resting-state functional magnetic resonance imaging (fMRI), has been shown to increase within the MTL in association with mild cognitive impairment [5], age-related cognitive decline [6], and AD [7]. Furthermore, a study involving healthy older individuals of African ancestry found that carriers of the ABCA7 rs115550680 G allele exhibited elevated MTL connectivity, which was linked to an increased risk of developing AD [8]. These findings suggest that MTL functional connectivity may serve as a promising biomarker for early detection and tracking the progression of AD.

However, static functional connectivity measures fall short in capturing the dynamic nature of brain network interactions. Time-varying functional connectivity may offer increased sensitivity to early neural alterations in AD [9]. One such dynamic measure is network flexibility, which quantifies how frequently a brain region reconfigures its connections over time [10]. Higher dynamic flexibility has been associated with improved memory performance in humans [11,12].

In preclinical stages of AD, alterations in MTL network dynamic flexibility may serve as early markers of pathology, correlate with cognitive decline, and reflect underlying neurodegeneration [12,13]. Therefore, this study aims to predict MTL dynamic flexibility, a key marker for cognitive adaptability and effective information processing.

### 2.1. Cognition

Older adults often exhibit reduced MTL dynamic flexibility in applying previously learned knowledge to new contexts, which contributes to difficulties in problem-solving [14]. Similarly, individuals in the preclinical stages of AD frequently show impairments in generalizing past experiences to novel situations [15]. These deficits are believed to reflect age-related changes in the MTL and prefrontal cortex, regions essential for integrating and transferring information across different contexts [16]. In this study, generalization, as measured by the Fish Task [17], is used as a cognitive predictor of MTL dynamic flexibility. Prior work has shown that performance on this task is associated with individual differences in MTL network dynamics, including flexibility, among older adults [11].

In addition, acquisition performance, which reflects the ability to learn initial stimulus-outcome associations, is also modeled as a predictor. Acquisition is strongly dependent on hippocampal and broader MTL integrity, particularly in tasks requiring the formation of arbitrary or relational associations [18]. Prior studies have shown that better acquisition performance is associated with greater hippocampal activation and connectivity [19]. Given that MTL dynamic flexibility reflects the brain’s capacity to dynamically reconfigure in response to new cognitive demands [11], individuals with stronger acquisition performance may exhibit more adaptive and resilient MTL network dynamics. Therefore, both acquisition and generalization are included as complementary predictors of individual variability in MTL dynamic flexibility, shedding light on how learning processes relate to neural adaptability.

### 2.2. Genetic

Genetic variation plays a critical role in Alzheimer’s disease, influencing both susceptibility and disease progression. In this study, two genetic biomarkers of AD—Apolipoprotein E (APOE) and ATP-binding cassette subfamily A member 7 (ABCA7)—measured via saliva samples, are examined as predictors of MTL dynamic flexibility. The ε4 isoform of APOE (APOE ε4) is the strongest known genetic risk factor for late-onset AD, associated with increased amyloid-beta (Aβ) accumulation, neuroinflammation, and synaptic dysfunction [20,21]. Conversely, the ε2 isoform (APOE ε2) appears to be protective, facilitating Aβ clearance and reducing tau pathology.

ABCA7, another lipid transporter gene linked to AD, has been particularly associated with elevated risk in individuals of African ancestry [8,22]. Research suggests that ABCA7 modulates the processing of amyloid precursor protein (APP), affecting the generation and clearance of Aβ peptides [23]. Loss-of-function mutations in ABCA7 can impair Aβ phagocytosis and promote amyloid plaque formation, one of the pathological hallmarks of AD.

Aβ accumulation begins years before clinical symptoms and is especially damaging to memory-related brain regions such as the hippocampus and surrounding MTL areas. Aβ disrupts synaptic signaling, promotes oxidative stress, and impairs both structural and functional connectivity [24,25]. These disruptions can interfere with the brain’s capacity for network dynamic flexibility, especially in the MTL. Given that dynamic flexibility reflects the ability to shift functional connections in response to changing demands, genetic variants that drive Aβ pathology may lead to reduced neural adaptability. Thus, APOE and ABCA7 genotypes are included as predictors to evaluate their impact on dynamic MTL network properties in preclinical AD.

### 2.3. Biochemical markers

The intracellular accumulation of hyperphosphorylated tau, forming neurofibrillary tangles, is another central hallmark of AD [26]. Elevated tau levels in cerebrospinal fluid (CSF) likely result from increased phosphorylation and neuronal secretion of tau in response to Aβ, marking early neurodegenerative changes [27]. Blood-based tau biomarkers such as plasma P-tau181 have been shown to correlate with both amyloid and tau PET imaging [28]. Recent findings indicate that P-tau217 outperforms P-tau181 in sensitivity and specificity, offering a more robust marker of AD pathology due to its stronger correlation with tau-PET [12,29]. Therefore, plasma P-tau217, measured through blood samples, is included as a biochemical predictor in this study.

### 2.4. Missing Values

A significant challenge in AD research is the prevalence of missing data, which is common in clinical and neuroimaging studies [30]. Missing data in AD studies can arise from several factors. Logistical constraints, such as requiring participants to make multiple visits to a research or clinical facility, can lead to incomplete datasets. Additionally, some participants may be unable to undergo MRI scans due to non-MRI-compatible implants, claustrophobia, or other medical contraindications. Cognitive assessments may also be incomplete due to participant fatigue, lack of motivation, or cognitive decline, preventing task completion. Furthermore, external factors such as human errors in data collection, budget limitations, evolving study protocols, and changes in funding sources can contribute to missing data, particularly in longitudinal studies.

Missing data mechanisms can be classified into three categories [31]: Missing Completely at Random (MCAR), Missing at Random (MAR), and Missing Not at Random (MNAR). In MCAR, data are missing for reasons unrelated to both observed and unobserved variables, meaning the likelihood of missingness is entirely random. For example, in a survey, some respondents may accidentally skip the question about their age due to a formatting issue in the survey software. In MAR, the probability of a missing value depends only on observed data, allowing for statistical estimation of the missing values. For instance, individuals with higher education may be more likely to withhold their income information in a study on earnings, not because of their actual income level but due to their educational background, an observed variable. Lastly, in MNAR, missingness depends on both observed and unobserved values. For example, employees who are dissatisfied with their jobs may be less likely to disclose their salary, as their unwillingness to report income is influenced not just by observable characteristics but also by the unobserved variable job satisfaction.

Addressing missing data is essential for maintaining the reliability and validity of research findings. Two common approaches for handling missing data are case deletion and imputation. Case deletion can be listwise, where an entire case is removed if it has any missing values, or pairwise, which minimizes data loss by excluding cases only from specific analyses requiring the missing data. Alternatively, imputation involves replacing missing values with predicted estimates based on observed data, helping to retain more information in the dataset. Choosing the appropriate method depends on the nature and extent of the missing data [32,33].

The high level of missingness, which is commonly encountered in medical and clinical datasets [34], highlights the importance of implementing appropriate missing data imputation strategies. First, using only the few complete cases for analysis would severely limit the reliability and generalizability of regression models due to the drastically reduced sample size. Second, although many cases are incomplete, they still contain substantial partial information that can contribute to the predictive power and robustness of statistical models. By leveraging imputation techniques, this incomplete yet valuable data can be retained and utilized, thereby improving the overall performance and validity of analyses.

To date, various techniques have been developed to impute missing values. Multiple Imputation by Chained Equations (MICE) [35], imputation using random forests (MissForest) [36] and Generative Adversarial Imputation Networks (GAIN) [37] are among the top-tier missing value imputation methods [38].

Each of these techniques has been effectively utilized in various medical and clinical studies to address high rates of missing data. For instance, in a study analyzing data from stroke patients, MICE missing value imputation enabled the development of a regression model to assess factors influencing the time from patient arrival to computed tomography (CT) imaging, a critical quality indicator in stroke care [39]. In another study, MissForest and MICE are used in two clinical datasets: a cirrhosis cohort (446 patients) and an inflammatory bowel disease cohort (395 patients) [40]. MissForest demonstrated the least imputation error for both continuous and categorical variables across varying frequencies of missingness. Additionally, predictive models utilizing MissForest imputed data exhibited the smallest prediction differences, underscoring its efficacy in maintaining predictive accuracy in clinical settings. Some studies show GAIN outperforming both MissForest and MICE in imputing missing data, especially at high missingness rates (50%) and for skewed continuous variables [41].

These techniques are explained in Section 3.1. Section 3.2 describes the regression models utilized to estimate MTL dynamic flexibility using available predictors.

## 3. Methods

This section describes in detail the methods for handling missing values and the regression models to estimate MTL dynamic flexibility using available predictors. Furthermore, metrics to assess the performance of each process are described below.

### 3.1. Handling Missing Values

This research employs four methods to address missing values, including case deletion, MICE, MissForest, and GAIN.

#### Case Deletion

Missing data were handled using the pairwise deletion approach [42], wherein only the available data for each set of variables were used in the analysis, allowing cases with some missing values to be partially retained. This approach ensures that only complete data for each specific analysis is included, helping to avoid potential bias introduced by imputation methods. However, this process results in fewer cases being available for fitting the regression model, which significantly reduces the model’s efficacy.

##### MICE

Multiple Imputation by Chained Equations (MICE) is a method for handling missing data by creating multiple complete datasets through iterative imputation [35]. Starting with the variable with the least number of missing values, each variable with missing values is modeled using the other variables in the dataset, and missing values are filled in based on these models. This process is repeated in cycles multiple times, generating several datasets that reflect the uncertainty of the missing data. The results from these datasets are then combined to produce final estimates. MiceForest [43] is utilized in this research, which is an upgraded implementation of MICE that employs gradient boosting trees, allowing for the robust imputation of binary variables. The utilized hyperparameters include a maximum of 50 iterations and 5 alternative imputations to generate the final aggregated predictions.

#### MissForest

MissForest is an imputation method that uses random forests to predict and fill in missing values [36]. It iteratively models each variable with missing data using the others as predictors, updating the imputed values until they stabilize. MissForest works well when the relationships between variables are nonlinear or complex and does not assume a specific data distribution. The process does not have any hyperparameters.

##### GAIN

Generative Adversarial Imputation Nets (GAIN) is a missing data imputation method that leverages the Generative Adversarial Network (GAN) framework [37]. It employs a generator network to impute the missing values conditioned on the observed data and a discriminator network to distinguish between the originally observed and the imputed values. To guide the generator towards learning the true data distribution, a hint mechanism provides the discriminator with partial information about the missingness pattern, encouraging the generator to produce more realistic imputations through an adversarial learning process. A batch size of 64, a hint rate of 0.90, an alpha of 60, and 1000 iterations are used as hyperparameters.

### 3.2. Regression Models

Five regression methods are employed in this study: Ridge Regression, k-Nearest Neighbors (k- NN), Support Vector Regression (SVR), regression trees, and Artificial Neural Networks (ANN). Hyperparameters for each model are selected using grid search with 3-fold cross-validation on the training dataset. Standard scaling, where predictor variables are standardized by removing the mean and scaling to unit variance, is applied as a preprocessing step for Ridge, k- NN, and SVR. In contrast, min-max scaling, where features are scaled to the [0, 1] range, is used for ANN.

#### Ridge Regression

Ridge Regression is a linear regression technique that penalizes correlated predictor variables, leading to a more stable and reliable regression model [44]. The penalty that is proportional to the square of the magnitude of the regression coefficients shrinks the correlated coefficients towards zero. The optimal value of the hyperparameter α that controls the shrinkage is selected through cross-validation.

#### k-NN

K-Nearest Neighbors (k-NN) regression predicts the target value of a data point by averaging the values of its k nearest neighbors, typically based on Euclidean distance [44]. This approach makes no assumptions about the data distribution and is easy to implement. The key hyperparameter is the number of neighbors k, selected via cross-validation.

##### SVR

Support Vector Regression (SVR) fits a regression function that approximates the data within a specified error margin (epsilon) while maintaining a model that is as simple and flat as possible [44]. Only data points that fall outside this margin, those with larger prediction errors, affect the model, which helps make SVR robust to outliers. To capture both linear and nonlinear relationships, SVR employs kernel functions such as linear, polynomial, or radial basis function (RBF), allowing it to adapt to various data patterns effectively. Key hyperparameters include the regularization parameter C, the margin width ε, and kernel-specific parameters (e.g., γ for RBF), which have been selected using cross-validation.

#### Regression Trees

Regression tree models are decision tree-based methods used to predict continuous target variables by recursively splitting data into subsets based on feature values to minimize prediction error [44]. Three tree-based ensemble methods are used: Bagging Trees, Random Forests, and Boosting Trees. Bagging Trees improves stability and accuracy by training multiple trees on different bootstrapped samples and averaging their predictions, reducing variance. Random Forests enhance bagging by also selecting a random subset of features at each split, increasing model diversity and further improving generalization. Boosting Trees builds trees sequentially, where each new tree corrects the errors of the previous ones, allowing the model to learn from its mistakes. Key hyperparameters used include the number of trees (1000), maximum samples (10 after case deletion, 50 after imputation), and learning rate (0.90) for boosting trees.

##### ANN

Artificial Neural Networks (ANNs) are computational models inspired by the human brain, designed to predict outcomes by learning complex patterns in data [44]. An ANN consists of layers of interconnected nodes (neurons), including an input layer, one or more hidden layers, and an output layer. Each neuron applies a weighted sum of inputs, followed by an activation function to capture nonlinear relationships. During training, the network adjusts its weights using algorithms like backpropagation to minimize the difference between predicted and actual values. Adam optimizer, batch size of 8, training epochs of 500, and one hidden layer with 5 neurons are used as model hyperparameters. No regularization, e.g., dropout, was applied, as initial testing showed no overfitting due to the relatively small model architecture.

### 3.3. Performance Assessment

After handling missing values, 5-fold cross-validation is used to evaluate the performance of the regression models on unseen data. In this approach, the data is randomly partitioned into five groups or folds. The regression models discussed in Section 3.2 are trained on four of these folds and then evaluated on the remaining one, the validation fold, using Mean Absolute Error (MAE):

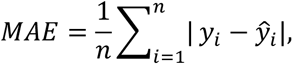

and Root Mean Squared Error (RMSE):

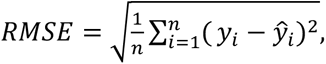

where n is the number of cases in the validation fold, *y_i_* is the observed target value, and *ŷ_i_* is its prediction using the regression model. This process is repeated five times, with each fold serving as the validation set exactly once. The performance metrics obtained from each of the five evaluations are then averaged to provide a more robust estimate of the model’s generalization ability. Finally, the 5-fold cross-validation process is repeated 25 times independently, and the average performance, including runtime, RMSE, and MAE, is reported to evaluate each combination of missing value handling methods and regression models. In this context, we considered MAE values below 0.1 to reflect high predictive accuracy, as they represent less than 10% absolute error relative to the observed range of MTL dynamic flexibility.

## 4. Materials

A total of 656 individuals of African ancestry (average age ± SD: 69.81 ±7.38 years; 80.6% female; average education years ± SD: 13.93 ± 2.29 years) were included in this study, recruited through the ongoing longitudinal Pathways to Healthy Aging and Brain Health Alliance study. This initiative investigates the relationships among genetics, cognition, health, and lifestyle factors in older individuals of African ancestry residing in the Greater Newark, NJ area. The study was conducted at Rutgers University–Newark and funded by the National Institute on Aging (NIA) under the following grant numbers: 1. Grant#1R01AG053961 (NIH/NIA) and 2.

Grant#R01AG078211. Further details on community engagement and recruitment strategies are available in Gluck et al. (2025) [45]. The study received ethical approval from the Rutgers University Institutional Review Board (IRB) and was conducted in accordance with the principles of the 1975 Helsinki Declaration.

All participants were fluent English speakers and provided written informed consent prior to participation. The initial screening was conducted over the phone. At this stage, individuals who self-reported a diagnosis of mild cognitive impairment (MCI) or dementia were excluded.

Additional exclusion criteria included the use of medications typically prescribed for cognitive impairments, a history of learning disabilities, colorblindness, self-reported excessive alcohol and/or drug use, or having undergone a medical procedure requiring general anesthesia within the past three months.

Blood samples were collected from all participants to measure biomarkers, including P-tau217. Additionally, most participants provided a saliva sample, which was used to assess APOE and ABCA7 genetic variants. Participants were also asked whether they consented to undergo an MRI session at the Rutgers University Brain Imaging Center (RUBIC); 34.15% of eligible participants agreed. From these sessions, Medial Temporal Lobe (MTL) dynamic flexibility was measured [11].

In detail, the MRI data were acquired using a 3T whole-body scanner equipped with a standard 32-channel head coil. Whole-brain acquisitions included a T1-weighted (T1w) magnetization-prepared rapid acquisition gradient echo (MP-RAGE) sequence [46], with parameters: voxel size = 1 × 1 × 1 mm³, FOV = 240 × 256 × 208 mm³, TR = 2300 ms, TE = 2.98 ms, and TI = 900 ms.

Additionally, a 2D multi-band echo-planar imaging (EPI) blood oxygen level-dependent (BOLD) sequence was used, as described in the ADNI4 dataset [47], with parameters: voxel size = 2.5 × 2.5 × 2.5 mm³, FOV = 220 × 220 × 160 mm³, TR = 607 ms, TE = 32.0 ms, and flip angle = 50°. The total scan time was approximately 7 minutes.

All participants completed either the Montreal Cognitive Assessment (MoCA) [48] or the Mini-Mental State Examination (MMSE) [49]. These two tests are widely used screening tools for detecting and monitoring cognitive impairments, such as those associated with Alzheimer’s disease, mild cognitive impairment (MCI), and other forms of dementia. To ensure consistency across assessments and enable direct comparisons, MMSE scores were converted to MoCA- equivalent scores using the method described by Fasnacht et al. [50].

Participants’ generalization and acquisition abilities were assessed using the Fish Task [17], which is freely available at https://github.com/Aging-and-Brain-Health-Alliance/GeneralizationTasks. This cognitive task is designed to evaluate learning, memory, and generalization, particularly in relation to associative learning and the integrity of the MTL dynamic flexibility, including structures such as the hippocampus. A summary of demographic information, along with summary statistics of the variables used in this study, is presented in Table 1.

**Table 1.** Summary of the variables in this study (N = 656).

| Variable | #valid cases (%) | # missing (%) | Range | Ave.(SD)/N (%) | Skewness |
| --- | --- | --- | --- | --- | --- |
| MTL dynamic flexibility | 224 (34.15%) | 432 (65.85%) | [0.09,0.99] | 0.42 (0.23) | 0.471 |
| Generalization | 284 (43.29%) | 372 (56.71%) | [0,1] | 0.60 (0.27) | -0.009 |
| Acquisition | 284 (43.29%) | 372 (56.71%) | [0.19,1] | 0.74 (0.14) | -0.523 |
| MoCA | 624 (95.12%) | 32 (4.88%) | [11,29] | 23.07 (3.46) | -0.548 |
| APOE* | 571 (87.04%) | 85 (12.96%) | 0,1 | high: 190 (33.3%) | 0.712 |
| ABCA7 50* | 608 (92.70%) | 48 (7.30%) | 0,1 | high: 306 (50.3%) | -0.013 |
| ABCA7 80* | 608 (92.70%) | 48 (7.30%) | 0,1 | high: 80 (13.2%) | 2.185 |
| Age (yrs) | 635 (96.80%) | 21 (3.20%) | [54,92] | 69.81 (7.38) | 0.344 |
| Sex | 649 (98.93%) | 7 (1.07%) | 0,1 | females: 529 (80.6%) | 1.627 |
| Education (yrs) | 646 (98.48%) | 10 (1.52%) | [6,22] | 13.93 (2.29) | 0.382 |
| P-tau217 | 217 (33.08%) | 439 (66.92%) | [0.06,1.87] | 0.34 (0.27) | 2.824 |

Note. For the variables highlighted with \*, we report the count and % for the high-risk cases.

## 5. Results

A total of 1,866 values were missing, representing approximately 25.86% of the dataset. However, when cases, i.e., individual participants with any missing values were removed, only 42 complete cases remained. This means that nearly 93.60% of the participants had at least one missing value. Using the available predictors, several state-of-the-art regression techniques were employed to model medial temporal lobe (MTL) dynamic flexibility. Table 2 and Figure 1 present the average validation MAE computed over 25 independent 5-fold cross-validation runs for each combination of missing data handling method and regression models.

**Figure 1.**
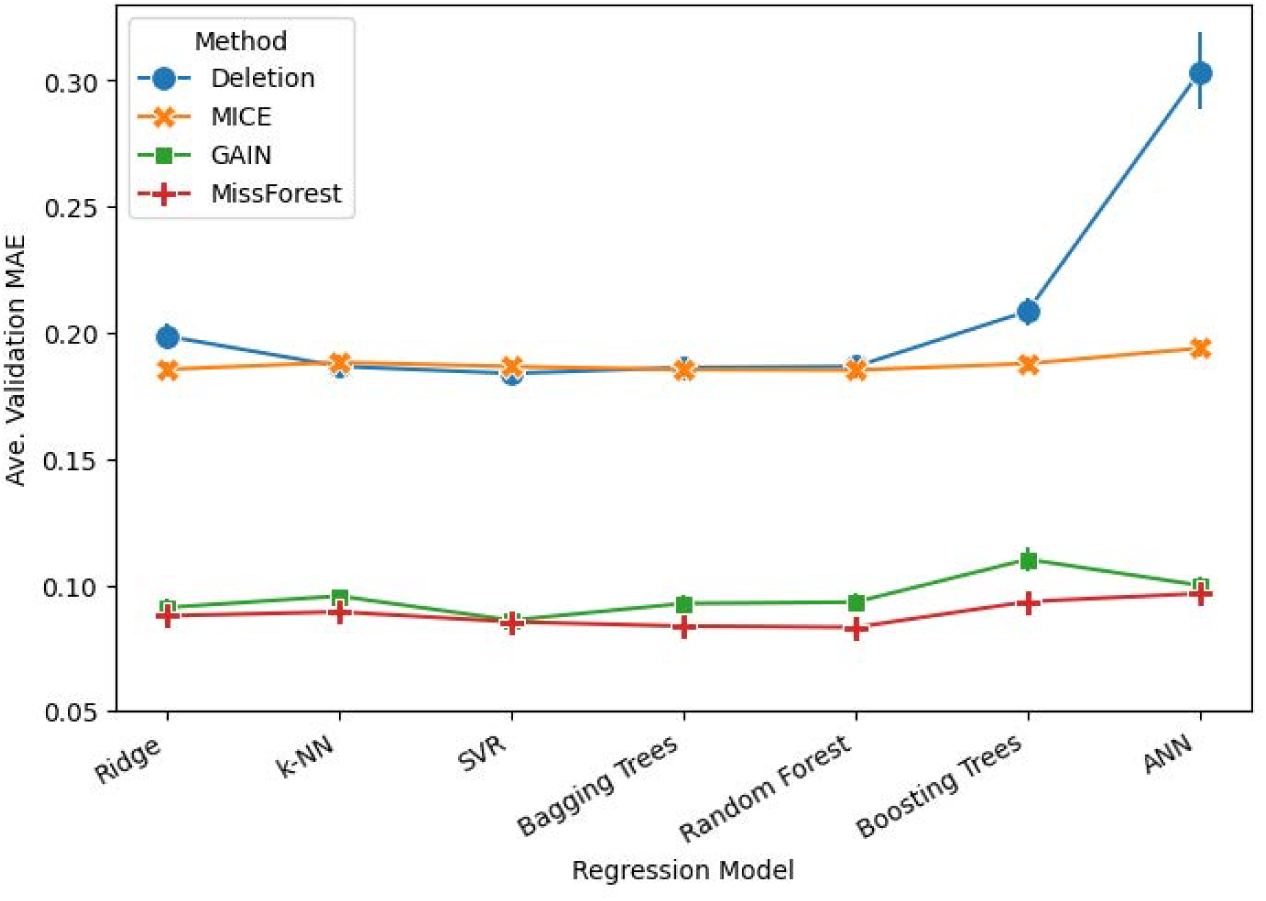
Average validation MAE and 95% error bars of regression models across different missing value handling methods.

**Table 2.** Average (SD) validation MAE from 25 independent runs for predicting MTL dynamic flexibility.

| Regression Model | Missing data Handling Technique |  |  |  | Average |
| --- | --- | --- | --- | --- | --- |
|  | Case Deletion | MICE | GAIN | MissForest |  |
| Ridge | 0.199 (0.0110) | 0.1854 (0.0058) | 0.091 (0.0067) | 0.088 (0.0005) | 0.141 (0.0060) |
| k-NN | 0.187 (0.0045) | 0.188 (0.0059) | 0.096 (0.0059) | 0.089 (0.0009) | 0.140 (0.0043) |
| SVR | 0.184 (0.0073) | 0.187 (0.0052) | 0.086 (0.0070) | 0.085 (0.0006) | 0.135 (0.0050) |
| Bagging Trees | 0.186 (0.0053) | 0.1854 (0.0059) | 0.093 (0.0063) | 0.084 (0.0005) | 0.137 (0.0045) |
| Random Forest | 0.187 (0.0049) | 0.1852 (0.0057) | 0.093 (0.0063) | 0.083 (0.0004) | 0.137 (0.0044) |
| Boosting Trees | 0.209 (0.0115) | 0.188 (0.0061) | 0.110 (0.0104) | 0.093 (0.0026) | 0.150 (0.0077) |
| ANN | 0.303 (0.0377) | 0.194 (0.0061) | 0.100 (0.0063) | 0.097 (0.0023) | 0.173 (0.0131) |
| Average | 0.208 (0.0117) | 0.187 (0.0058) | 0.096 (0.0070) | 0.088 (0.0011) | 0.145 (0.0064) |
| Note. Details of runs are provided in Table 1 of supplementary materials. |  |  |  |  |  |

**Table 3.** Average (SD) of runtime (seconds) from 25 independent runs.

| Regression Model | Missing data Imputation Technique |  |  |  |
| --- | --- | --- | --- | --- |
|  | MICE | GAIN | MissForest | Average |
| k-NN | 0.34 (0.005) | 0.35 (0.004) | 0.34 (0.004) | 0.34 (0.005) |
| SVR | 0.17 (0.003) | 0.18 (0.005) | 0.18 (0.003) | 0.18 (0.004) |
| Bagging Trees | 2.95 (0.152) | 2.02 (0.055) | 3.77 (0.657) | 2.91 (0.288) |
| Random Forest | 1.59 (0.013) | 1.62 (0.022) | 1.63 (0.010) | 1.61 (0.015) |
| Boosting Trees | 0.98 (0.008) | 1.00 (0.010) | 0.99 (0.007) | 0.99 (0.008) |
| ANN | 0.06 (0.035) | 0.04 (0.015) | 0.09 (0.037) | 0.06 (0.029) |

Note. Details of runs are provided in Table 1 of supplementary materials.

The first phase of the research focused on the 42 complete cases, which represent only 6.40% of the participants. Among the models evaluated, SVR achieved the best performance, with an average (SD) MAE of 0.184 (0.0073), indicating that, on average, predictions deviated from the true values by approximately 0.184 ± 0.0073 units. Given that the target variable ranges from 0.09 to 0.99, this level of error is relatively high and suggests limited predictive accuracy. The ANN model produced the highest average (SD) MAE of 0.303 (0.0377), which is not unexpected given that neural networks typically require large datasets to build reliable predictive models. The average (SD) MAE across all regression models was 0.208 (0.0117), further underscoring the limitations of working with such a small subset of complete cases.

In the second phase of the study, the three imputation techniques described in section 3.1 were applied to estimate missing values. Following imputation, MTL dynamic flexibility was modeled using the same set of regression techniques within a 5-fold cross-validation framework. The average (SD) validation MAEs from 25 independent runs are reported in Table 2 and visualized in Figure 1.

As the MAE values were approximately normally distributed at the 99% significance level (Shapiro–Wilk test, p > 0.029 for all) but demonstrated significant heteroskedasticity in variance (Bartlett’s and Levene’s tests, p < 0.001), the robust Scheirer–Ray–Hare ANOVA was employed to assess the differences in model performance across missing data handling methods. The analysis revealed a statistically significant effect of imputation method on prediction performance (p < 0.001), as measured by average validation MAE. All imputation methods substantially outperformed case deletion. Among them, MissForest was the most effective, reducing the average MAE by 57.4%, followed by GAIN (54%) and MICE (9.7%). However, MICE did not outperform case deletion over all regression models, as its performance was statistically the same in k-nearest neighbors (k-NN), support vector regression (SVR), bagging trees, and random forest.

Furthermore, support vector regression (SVR) consistently emerged as the best-performing regression model (p < 0.001), achieving the lowest average validation MAE across all missing value handling methods. Other models, including random forest, bagging trees, k-NN, and ridge regression, also demonstrated competitive performance. In contrast, boosting trees and ANNs yielded slightly higher error rates. Overall, the MissForest imputation paired with random forest regression achieved the lowest prediction error (MAE ± SD: 0.083 ± 0.0004), making it the most accurate and reliable strategy for modeling MTL dynamic flexibility under high missingness conditions. This MAE represents less than 10% absolute deviation from observed MTL dynamic flexibility scores, underscoring the model’s practical utility. Compared to the best-performing case deletion method, this represents a 54.7% improvement in predictive accuracy.

Figure 2 presents the imputation times for each method. Among them, GAIN was the fastest, requiring an average (SD) of 3.17 (0.115) seconds to impute all 1,866 missing values. MissForest followed with an average (SD) time of 3.41 (0.069) seconds, while MICE took significantly longer at 5.97 (0.294) seconds.

**Figure 2.**
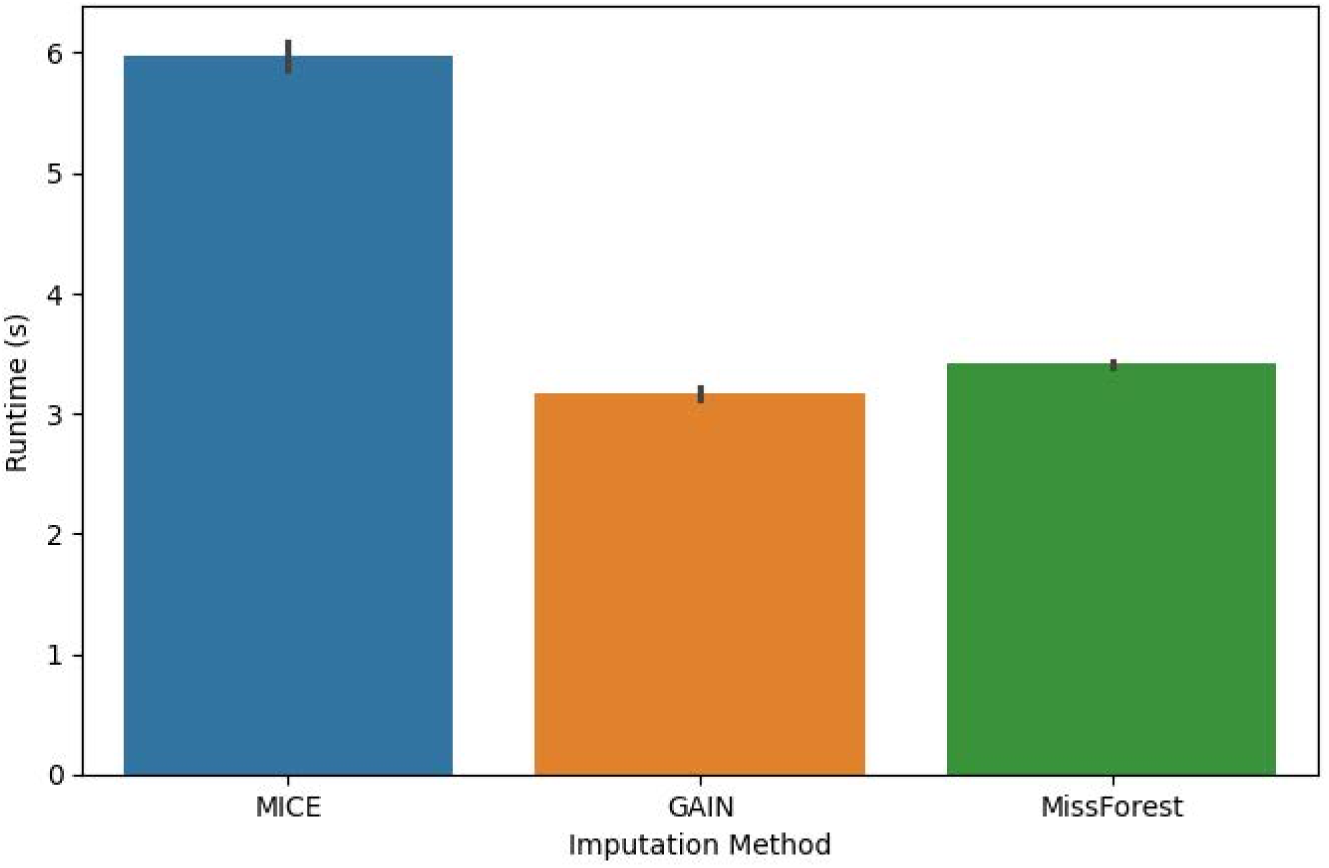
Average imputation time.

**Error! Reference source not found.** shows a significant variation in the runtime of regression models following imputation. Boosting trees emerged as the fastest, completing model fitting with five-fold cross-validation in just 0.06 seconds, followed by k-NN (0.18 s), ridge regression (0.34 s), random forest (0.99 s), bagging trees (1.61 s), and SVR (2.91 s). Artificial neural networks (ANN), implemented using TensorFlow 2, were by far the slowest, requiring 43.37 seconds on average. Notably, ANN models not only exhibited the longest runtimes but also delivered the poorest predictive performance among all evaluated regression methods.

Table 4 presents the summary statistics of the variables following missing value imputation by MissForest. The distributions of the variables remained consistent with those of the original data, preserving both the range and the central tendencies. The means and standard deviations of the imputed datasets closely matched the original values, indicating that the imputation did not introduce substantial bias or distortion. This preservation of key statistical properties is essential for maintaining the interpretability and comparability of the variables, particularly when they are used as predictors in regression models. It also suggests that the imputation technique effectively filled in missing values without significantly altering the overall structure of the dataset.

**Table 4.** Summary of the variables after missing value imputation using MissForest (N = 656)

| Variable | #vsolid cases (%) | Range | Ave. (SD)/N (%) | Skewness |
| --- | --- | --- | --- | --- |
| MTL dynamic flexibility | 656 (100%) | [0.09,0.99] | 0.43 (0.14) | 0.560 |
| Generalization | 656 (100%) | [0,1] | 0.59 (0.18) | 0.143 |
| Acquisition | 656 (100%) | [0.19,1] | 0.75 (0.09) | -0.960 |
| MoCA | 656 (100%) | [11,29] | 23.05 (3.38) | -0.546 |
| APOE* | 656 (100%) | 0,1 | high: 191 (33.3%) | 0.922 |
| ABCA7 50* | 656 (100%) | 0,1 | high: 354 (54.0%) | -0.159 |
| ABCA7 80* | 656 (100%) | 0,1 | high: 80 (12.2%) | 2.316 |
| Age (yrs) | 656 (100%) | [54,92] | 69.81 (7.27) | 0.351 |
| Sex | 656 (100%) | 0,1 | females: 536 (81.7%) | 1.644 |
| Education (yrs) | 656 (100%) | [6,22] | 13.94 (2.28) | 0.382 |
| P-tau217 | 656 (100%) | [0.06,1.87] | 0.30 (0.16) | 4.785 |

Note. For the variables highlighted with \*, we report the count and % for the high-risk cases.

## 6. Conclusions

This study underscores the critical importance of handling missing data in clinical datasets, particularly those related to Alzheimer’s disease research. With nearly 94% of cases exhibiting some degree of missingness, relying solely on complete cases would have drastically reduced the sample size and compromised model reliability. Our findings demonstrate that imputation not only preserves valuable partial data but also significantly enhances predictive performance.

Among the evaluated methods, the combination of missing value imputation using random forests (MissForest) and support vector regression (SVR) yielded the most accurate and efficient results, reducing prediction error by 54.7% compared to case deletion. These results are consistent with prior studies showing that MissForest often outperforms traditional imputation methods like MICE in both continuous and categorical data scenarios, particularly when handling complex, nonlinear relationships in biomedical datasets [36,40].

MissForest also proved to be computationally efficient, further supporting its practical utility in clinical research. In contrast, neural networks performed poorly and were computationally expensive, underscoring the importance of aligning model complexity with dataset characteristics, especially in studies with limited sample sizes or computing resources. This observation is supported by existing literature showing that neural networks often require large, complete datasets to reach optimal performance and may not be the most suitable choice for small or highly sparse clinical data [51].

Overall, our results advocate for the adoption of robust imputation strategies, particularly MissForest, paired with Random Forest regression to improve MTL dynamic flexibility prediction accuracy and generalizability in the presence of substantial missing data. These findings align with previous reports emphasizing the synergy between robust imputation and ensemble methods in clinical prediction tasks [52,53]. While the models in this study demonstrated strong performance within a cohort of older individuals of African ancestry, a group often underrepresented in neuroscience research, their generalizability to other populations and settings remains to be tested.

Although both MAE and RMSE were computed throughout the study, only MAE values are reported in the main text due to their more intuitive interpretability. Nonetheless, RMSE-based results mirrored the trends observed with MAE, as shown in Supplementary Table 1, which presents the mean (SD) validation RMSE from 25 independent runs for predicting MTL dynamic flexibility.

Not all imputation methods are equally effective, and their suitability depends heavily on data structure and the underlying mechanism of missingness. Despite its widespread use in clinical research, MICE underperformed in this analysis, likely due to its limited ability to model complex, nonlinear, and high-dimensional relationships, even when implemented via gradient-boosted trees (MiceForest). This finding is consistent with prior work showing that MICE may falter under conditions of high multicollinearity, nonlinearity, or interactions among predictors [52,54]. Therefore, selecting an imputation method requires a careful evaluation of the dataset’s properties and the modeling objectives to preserve validity and interpretability. Future research is needed to assess the longitudinal utility of these models, particularly whether early estimates of MTL dynamic flexibility, obtained through imputation and regression, can reliably predict subsequent cognitive decline or conversion to AD.

## Abbreviations

AD: Alzheimer’s Disease
MTL: Medial Temporal Lobe
MICE: Multiple Imputation by Chain Equations
MissForest: Missing value imputation using random Forests
GAIN: Generative Adversarial Imputation Networks
MRI: Magnetic resonance imaging
CT: Computed Tomography
MoCA: Montreal Cognitive Assessment
MMSE: Mini-Mental State Examination
APOE: Apolipoprotein E
ABCA7: Adenosine Triphosphate Binding Cassette Subfamily A Member 7
MCAR: missing completely at random
MAE: Mean Absolute Error
RMSE: Root Mean Squared Error
SD: Standard Deviation
k-NN: k-Nearest Neighbors
SVR: Support Vector Regression
ANN: Artificial Neural Networks
EPI: Echo-Planar Imaging
BOLD: Blood Oxygen Level-Dependent.

## Author contributions

All authors contributed to the study and approved the final version.

## Acknowledgments

The authors would like to acknowledge the support from NIA and NIH for funding. We also appreciate the help and efforts from our community engagement team.

